# MERIT: controlling Monte-Carlo error rate in large-scale Monte-Carlo hypothesis testing

**DOI:** 10.1101/2022.01.15.476485

**Authors:** Yunxiao Li, Yi-Juan Hu, Glen A. Satten

## Abstract

The use of Monte-Carlo (MC) *p*-values when testing the significance of a large number of hypotheses is now commonplace. In large-scale hypothesis testing, we will typically encounter at least *some p*-values near the threshold of significance, which require a larger number of MC replicates than *p*-values that are far from the threshold. As a result, the list of detections can vary when different MC replicates are used, resulting in lack of reproducibility. The method of Gandy and Hahn (GH) (2014; 2016; 2017) is the only method that has directly addressed this problem, defining a Monte-Carlo error rate (MCER) to be the probability that any decisions on accepting or rejecting a hypothesis based on MC *p*-values are different from decisions based on *ideal p*-values, and then making decisions that control the MCER. Unfortunately, GH is frequently very conservative, often making no rejections at all and leaving a large number of hypotheses “undecided”. In this article, we propose MERIT, a method for large-scale MC hypothesis testing that also controls the MCER but is more statistically efficient than the GH method. Through extensive simulation studies, we demonstrated that MERIT controlled the MCER and substantially improved the sensitivity and specificity of detections compared to GH. We also illustrated our method by an analysis of gene expression data from a prostate cancer study.

## 1. Introduction

Modern scientific studies often require testing a large number of hypotheses; in particular, modern biological and biomedical studies of -omics such as genomics, proteomics, and metabolomics typically test a large number (≫ 100) of hypotheses at a time. In these studies, *p*-values for testing individual hypotheses are first obtained and a procedure that corrects for multiplicity, such as the procedures of Benjamini and Hochberg (BH) (1995), Bonferroni (1936), or Holm (1979), is then applied to make decisions on the hypotheses. When the *p-* values cannot be calculated analytically, they are frequently obtained by resampling methods such as permutation or bootstrap. For ease of exposition, hereafter we generically refer to any resampling-based method as a Monte-Carlo (MC) method, and refer to individual resamples as MC replicates.

The recommended form of an MC *p*-value is 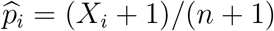 (Davison and Hinkley, 1997; Manly, 2006; Phipson and Smyth, 2010) for the *i*th hypothesis, where *n* is the number of MC replicates (i.e., MC sample size) and *X_i_* is the number of “exceedances”, i.e., when the statistic based on an MC replicate “exceeds” the observed test statistic. When using a finite *n*, there will be a difference between the MC *p*-value and the *ideal p*-value that is based either on analytical methods or exhaustive resampling of replicates. It is also well known that the MC sample size required to test a hypothesis having a *p*-value near a threshold of significance is much larger than that required for a hypothesis having a *p*-value that is far from the threshold. What is less appreciated is that, when testing a large number of hypotheses, it is very likely that at least *some* hypotheses have ideal *p*-values near whatever threshold of significance is used. As a result, there can be noticeable variability in the list of hypotheses rejected by an MC test procedure in different runs that use different MC replicates. The list can also differ from that based on the ideal *p*-values. This is true whether the test procedure adopts a fixed stopping criterion, i.e., mandating a fixed MC sample size for all hypotheses, or a sequential stopping criterion (Sandve et al., 2011), i.e., allowing some tests to stop early after collecting sufficient information about its decision.

To date, only Gandy and Hahn (GH) (2014; 2016; 2017) have *directly* addressed this problem. They developed a method that can be coupled with any MC test method that has a sequential stopping criterion. Their method makes decisions on multiple hypotheses by controlling the family-wise Monte-Carlo error rate (MCER), which is the probability that any of their decisions on accepting or rejecting a hypothesis are different from what would have been obtained from the ideal *p*-values. It works in a sequential manner, giving a decision among “rejected”, “accepted”, and “undecided” to each test after each iteration and continuing sampling only for hypotheses that are “undecided”.

To be more specific, denote the *m* null hypotheses by *H*_1,0_, *H*_2,0_,…, *H*_*m*,0_, the ideal *p*-values by 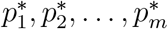, and the ideal decisions (between “rejected” and “accepted”) made by the BH procedure that is applied to the ideal *p*-values by 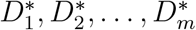. Denote the decisions (among “rejected”, “accepted”, and “undecided”) made by an MC test procedure by 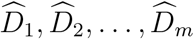. Following GH, we define the total number of MC errors to be the sum of all disagreements between the 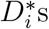 and 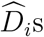 among hypotheses that are determined as “rejected” or “accepted” by the MC test procedure:

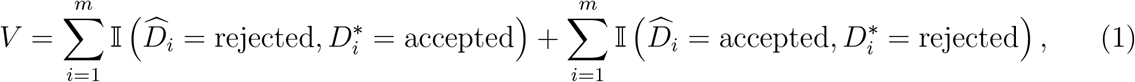

where 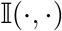 is an indicator function. Controlling the family-wise MCER at level *α*, i.e., Pr(*V* ≥ 1) ≤ *α*, assures that, among those tests that the MC test procedure actually decides, the probability that we reach a conclusion that is different from the decision we would make if we had the ideal *p*-values is less than *α*. Note that there is no error control on “undecided” hypotheses 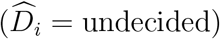, which would require more MC replicates to reach a decision.

At each iteration of the MC test procedure, the GH method forms a two-sided Robbins-Lai confidence interval (CI) (Darling and Robbins, 1968; Lai et al., 1976) for each ideal *p*-value that maintains a joint coverage over all iterations (i.e., accounting for the possible differences in MC sample size for each test); the level of each CI is determined by using the Bonferroni correction. Then, the GH method applies the BH procedure to the collection of upper CI limits and, separately, to the collection of lower CI limits. Those tests for which both lower and upper limits are rejected are determined as “rejected”, those for which both limits are accepted are determined as “accepted”; the remaining tests are determined as “undecided”. The GH method controls MCER at every iteration. However, it is extremely conservative due to the use of the Bonferroni correction, and thus tends to make no rejections at all when the MC sample size is not very large.

While the sequential stopping criterion has advantages (e.g., it can greatly reduce the MC sample size for tests that are either highly insignificant or highly significant), use of a fixed, pre-determined MC sample size is more common. Further, in many instances there may be little or no computational advantage to stopping some tests early, e.g., when the cost of evaluating the stopping criterion exceeds the cost of calculating the test statistics. Under these circumstances, the cost of satisfying an MCER at each iteration may not be justified. For these reasons, we now restrict our attention to MC procedures with fixed MC sample sizes.

In this article, we propose a method, called MERIT (<u>M</u>onte-Carlo <u>E</u>rror <u>R</u>ate control <u>I</u>n large-scale MC hypothesis <u>T</u>esting), that makes decisions on the hypotheses by controlling the same family-wise MCER as the GH method, but it works for MC test procedures with a fixed stopping criterion. We aim to maximize detection efficiency by minimizing the number of “undecided” hypotheses at a given MC sample size or by making conclusive decisions for all hypotheses with fewer MC replicates. We present our method in Section 2, examine its properties through simulation studies in Section 3, and apply it to a study of prostrate cancer with gene expression data in Section 4. Because one immediate advantage of using a fixed MC sample size is that the Robbins-Lai CI required for the sequential GH approach is much wider than any single-stage CI (Coe and Tamhane, 1993), in Section 2 we also develop a fixed-MC-sample-size version of GH that uses a single-stage CI, to allow for a more fair comparison in Sections 3 and 4. Finally, we conclude with a discussion in Section 5.

## 2. Methods

Suppose that a fixed number, *n*, of MC replicates have been collected for testing all *m* hypotheses. For testing the *i*th hypothesis *H*_*i*,0_, let *s_i_* be the observed test statistic and *S_ij_* be the test statistic calculated from the *j*th MC replicate. The input of our method is the matrix *I* = {*I_ij_*}_*m*×*n*_, where 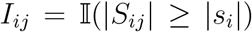 is the exceedance indicator corresponding to a symmetric, two-sided statistic (note that *I_ij_* can be redefined for one-sided tests or tests with asymmetric null distributions). We frequently use the total number of exceedances for each test, 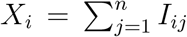. Because the replicates are sampled independently from the null distribution, we have 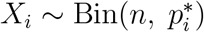, which is a binomial variable with *n* trials and “success” rate being the ideal *p*-value 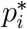.

We first transform the problem of controlling the MCER into a problem of controlling the family-wise error rate (FWER) in testing a set of new but related hypotheses. Then, because step-wise procedures (where the criterion for rejecting hypotheses becomes less stringent in subsequent steps in case some hypotheses have been rejected in an earlier step) such as Holm’s (1979) are typically more powerful than single-step procedures such as the Bonferroni correction, we develop a step-wise procedure for testing the new hypotheses following the framework of Romano and Wolf (2005; 2018). Because the Romano and Wolf framework for one-sided hypotheses (Romano and Wolf, 2018) can have additional power gains by adopting Hansen’s adjustment (Hansen, 2005), we focus on one-sided hypotheses by converting two-sided hypotheses into pairs of one-sided hypotheses.

Corresponding to the pairs of one-side hypotheses, we partition the total MC errors in (1) into the type-I and type-II MC errors, which are defined by the first and second terms in (1) and denoted by *V*^I^ and *V*^II^, respectively. We develop a procedure for testing a set of one-sided hypotheses, which makes decisions on the original hypotheses between “rejected” and “non-rejected” while controlling the family-wise type-I MCER Pr(*V*^I^ ≥ 1) at *α*_1_; we develop another procedure for testing another set of one-sided hypotheses, which makes decisions on the original hypotheses between “accepted” and “non-accepted” while controlling the familywise type-II MCER Pr(*V*^II^ ≥ 1) at *α*_2_. After combining their results, we can determine each original hypothesis as “rejected”, “accepted”, or possibly “undecided”, while controlling the family-wise MCER at *α*_1_ + *α*_2_, because Pr(*V* ≥ 1) ≤ Pr(*V*^I^ ≥ 1) + Pr(*V*^II^ ≥ 1). In what follows, we omit the term “family-wise” for simplicity. In Sections 2.1–2.3, we present our first procedure. In Section 2.4, we obtain the second procedure by modifying the first procedure. In Section 2.5, we show how to combine results of the two procedures.

### 2.1 A procedure that rejects original hypotheses while controlling the type-I MCER

Suppose that the BH procedure with nominal false-discovery rate (FDR) *ϕ* is applied to the ideal *p*-values, which are sorted in an ascending order 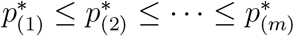. If *k* is the largest integer such that 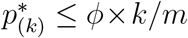, then the hypotheses corresponding to 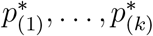 are rejected and the remaining hypotheses are not. Thus, we call *τ*\* ≡ *ϕ* × *k*/*m* the *BH cutoff* for the ideal *p*-value, which separates the ideal *p*-values for rejected and non-rejected hypotheses.

If *τ*\* is known *a priori*, the result of the above BH procedure can be viewed as the truth of the following hypotheses:

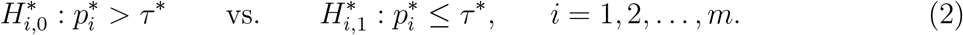

Recall that a type-I MC error occurs when a hypothesis 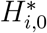 is true (i.e., not rejected using the ideal *p*-value) but rejected using the MC *p*-value; this would happen when the MC *p*-value is smaller than its corresponding ideal *p*-value. Then, a test procedure based on MC *p*-values that controls the FWER for testing 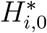 at *α*_1_ would control the type-I MCER at *α*_1_.

In reality, *τ*\* is unknown since the ideal *p*-values are unknown. A reasonable strategy to control type-I MCER would be to choose some threshold *τ* satisfying *τ* < *τ*\*; this is intuitive, since, as we make *τ* smaller, we require smaller MC *p*-values to reject a hypothesis, and smaller MC *p*-values correspond to increased confidence in any rejections we make. Thus, we divide the problem of testing hypotheses in (2) into two sub-problems and develop a two-step procedure. In the first step, we find a lower limit *τ_l_* for *τ*\* that satisfies *τ_l_* < *τ*\* with a probability of at least 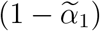, where 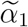 is chosen such that 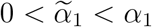. In the second step, we consider the revised hypotheses, treating *τ_ℓ_* as a constant:

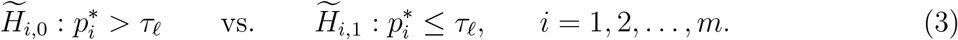

We develop a test procedure that tests (3) while controlling the FWER at level 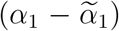.

Note that the same decision on 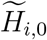 is transferred to 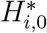 and *H*_*i*,0_. Then, by the relationship

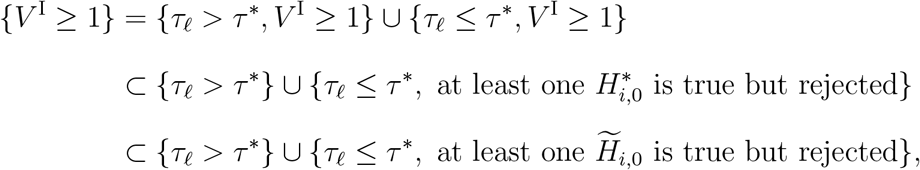

the type-I MCER Pr(*V*^I^ ≥ 1) is always bounded by the sum of Pr(*τ_ℓ_* > *τ*\*) and the FWER of the test procedure in testing (3), which is 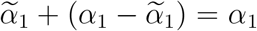. As the default, we choose 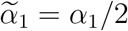, i.e., partitioning the type-I MCER equally into the two sub-problems. We show in Supplemental Materials S1 that the equal partition scheme generally yields the highest power.

### 2.2 Step 1: construct a 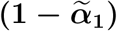-level one-sided CI [*τ_ℓ_*, 1] for *τ*\*

We now turn our attention to constructing a one-sided CI [*τ_ℓ_*, 1] for *τ*\*. Note that, for any set of *p*-values, there is a corresponding BH cutoff; thus, we find the lower limit *τ_ℓ_* for *τ*\* from a set of *p*-values that are “lower limits” for the ideal *p*-values 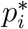. To this aim, we consider the following sets of *p*-values, with each set indexed by a positive continuous value *c*:

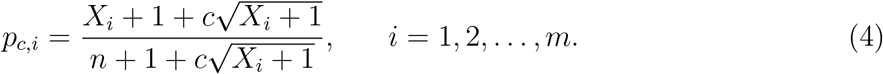

Recall that the MC *p*-value 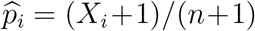 provides a consistent estimate of 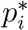. The term 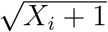 in (4) is an approximation to 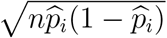, which is the estimated standard error of *X_i_* + 1; the term 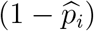 is omitted because only the small *p*-values (in the neighborhood of *τ*\*) are of interest. As *c* goes to 0, *p_c,i_* converges to 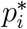. Because 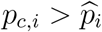 as long as *c* > 0 and *X_i_* < *n*, when *c* is sufficiently large, *p_c,i_* is asymptotically larger than 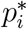, at least for small 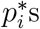.

Let *F*\*(*t*) and *F_c_*(*t*) be the empirical distribution functions (EDF) of 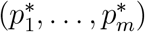 and (*p*_*c*,1_,…, *p_c,m_*), respectively, and *τ*\* and *τ_c_* be their BH cutoffs. Storey et al. (2004) gave a general result that, a BH cutoff *τ* can also be obtained from an EDF *F*(*t*) of *p*-values by the formula *τ* = max{*t*: *F*(*t*) ≥ *t*/*ϕ*, *t* ∈ [0,1]}, which means that *τ* is the intersection of *F*(*t*) and the straight line *t*/*ϕ*, as illustrated in Figure 1. When *t*/*ϕ* is constrained to [0,1] as *F*(*t*), *t* must be constrained to [0, *ϕ*]. Then we see from Figure 1 (right) that the intersection *τ* must lie in [0, *ϕ*] and is fully determined by the EDF of *p*-values in the range of [0, *ϕ*]. If *F_c_*(*t*) is always no greater than *F*\*(*t*) over [0, *ϕ*], as shown in Figure 1 (left), we must have *τ_c_* ≤ *τ*\*. It follows that

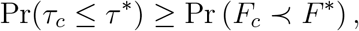

in which we use 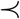 to denote that one function is always no greater than another function for *t* ∈ [0, *ϕ*]. Note that *τ_c_* and *F_c_* are random because *p_c,i_* involves *X_i_* that is based on MC replicates; *τ*\* and *F*\* are constant because we condition on the observed data. To construct a 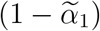-level one-sided CI [*τ_ℓ_*, 1] for *τ*\*, we are interested in *c* that satisfies

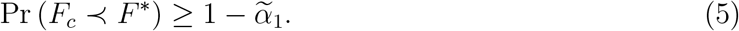

**Figure 1:**
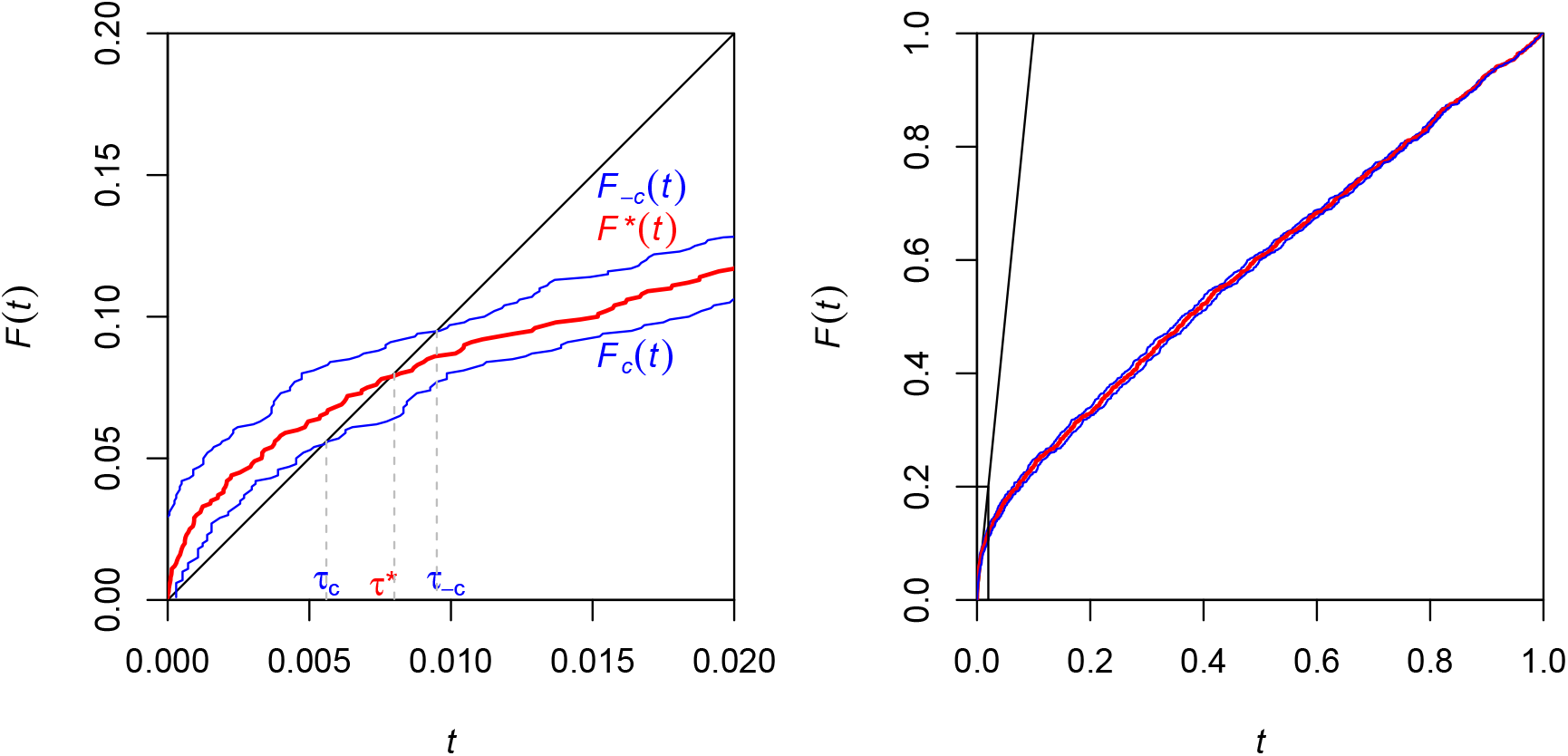
An example of *F*\*(*t*), *F_c_*(*t*), and *F*_−*c*_(*t*) and their corresponding BH cutoffs *τ*\*, *τ_c_*, and *τ*_−*c*_. The black straight line is *t*/*ϕ*, where *ϕ* = 10%. The left plot is a zoom-in view of the bottom left corner of the right plot.

The probability on the left of (5) increases as *c* increases (because the increase of *p_c,i_* leads to the decrease of *F_c_*(*t*) at a given *t*). To make the CI for *τ*\* the tightest, we wish to find the smallest *c* that satisfies (5), denoted by *c_ℓ_*. Finally, we set the lower limit *τ_ℓ_* to *τ_c_ℓ__*, which is the BH cutoff corresponding to the set of *p*-values in (4) indexed by *c_ℓ_*.

It is difficult to find the smallest *c* that satisfies (5) analytically because *F*\* is unknown. Thus, we propose to obtain *c_ℓ_* by bootstrap resampling. In the *b*-th bootstrap, we sample *n* columns from the input matrix *I* with replacement to form the bootstrap matrix *I*^(*b*)^ and sum up the rows of *I*^(*b*)^ to obtain the corresponding numbers of exceedances 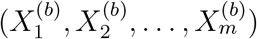.

The number of bootstrap replicates *B* needs to be large, e.g., *B* = 10^4^. Define the bootstrap *p*-values indexed by *c* as

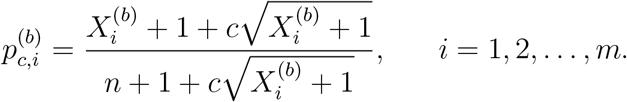

Let 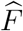 and 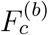 be the EDF for 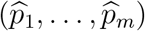 and 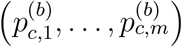, respectively. Then we wish to find *c_ℓ_* to be the smallest *c* that satisfies

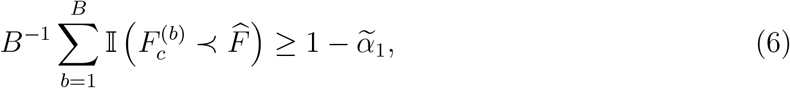

which is an empirical version of (5). In fact, it is still difficult to directly find the smallest *c* that satisfies (6). Alternatively, we find, for every *b*, the smallest *c* that guarantees 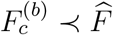, denoted by 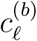, and then set *c_ℓ_* to be the 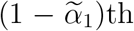 quantile of 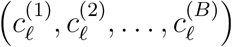. This strategy ensures that 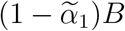 indicator functions in (6) take the value of 1 when evaluated at *c* = *c_ℓ_*. The procedure here is summarized in Step 1 of Algorithm 1.

#### Algorithm 1 The procedure that rejects original hypotheses while controlling type-I MCER

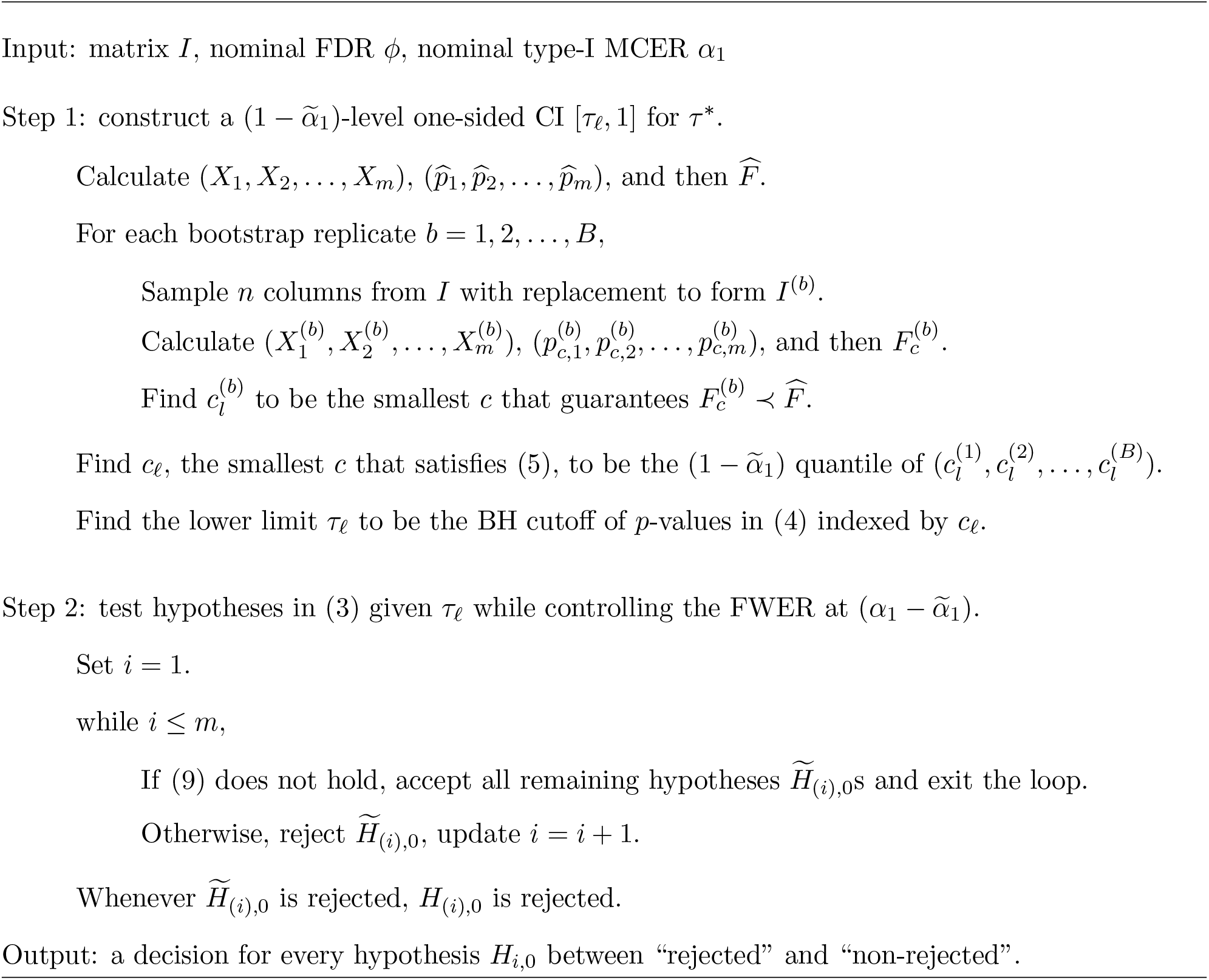

### 2.3 Step 2: test hypotheses in (3) given *τ_ℓ_* while controlling the FWER at 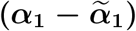

We use *X*_1_, *X*_2_,…, *X_m_* defined earlier as the test statistics and let *x*_1_, *x*_2_,…, *x_m_* denote their observed values. Let *x*_(1)_ ≤ *x*_(2)_ ≤… ≤ *x*_(m)_ be the ordered observed values, and 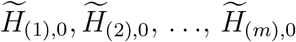 be the corresponding hypotheses.

We start by testing the global null 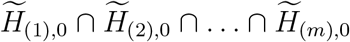, which means that all 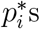 are greater than *τ_ℓ_*. We use *x*_(1)_ as the test statistic for this test. Romano and Wolf (2005; 2018) proposed to calculate the *p*-value under the global null in this form:

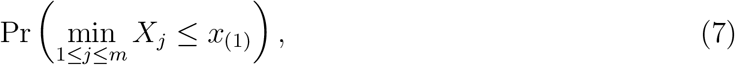

where each *X_j_* is a binomial random variable with *n* trials and some “success” rate *θ_j_* that is under the null and determined below. It is likely that *X*_1_, *X*_2_,…, *X_m_* have some dependence structure in many applications; Romano and Wolf proposed to calculate (7) via bootstrap resampling that preserves the dependence structure. To avoid bootstrap resampling, we use the Bonferroni inequality to obtain an upper bound for (7), namely

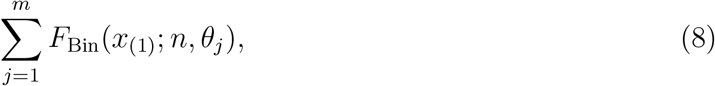

where *F*_Bin_(*x*; *n*, *θ*) denotes Pr(*X* ≤ *x*) for *X* ~ Bin(*n*, *θ*). Expression (8) can be calculated analytically, although this simplification comes at the cost of losing some statistical power compared to using (7) directly. Note that the upper bound in (8) is valid under any dependence structure among the *m* tests.

Now we consider the problem of determining *θ_j_*s. In the worst case when all 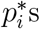 are close to the boundary *τ_ℓ_*, we should choose *θ_j_* = *τ_ℓ_* for all *j*s. Hansen (2005) noted that it is possible to gain more power by making adjustments for null hypotheses that are “deep in the null”, i.e., 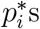 that are not near the boundary *τ_ℓ_* but instead satisfy 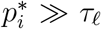. In the presence of a large number of tests, we expect many 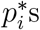 are “deep in the null”. Intuitively, the hypotheses associated with very large 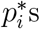 can be removed and the “effective” number of tests that we need to adjust for can be considerably reduced, leading to improved power. Ideally, we should choose 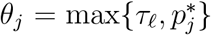. However, 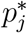 is unknown. Hansen (2005) proposed an estimator for 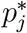 that leads to the choice

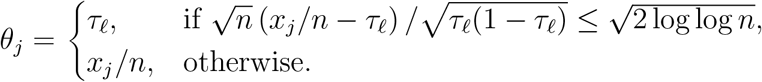

Therefore, if the statistic *x_j_* is sufficiently large, *θ_j_* is set to the sample-based estimator *x_j_*/*n*, and otherwise it is left unchanged at *τ_ℓ_*.

Using this *θ_j_*, if

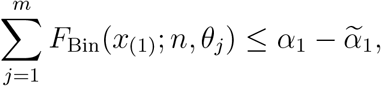

we reject 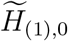 and move to the next joint null 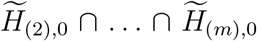 that excludes 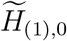; otherwise, we stop and declare that none of the hypotheses should be rejected. In general, for testing the joint null 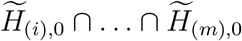 for any *i*, we use *x*_(*i*)_ as the test statistic. If

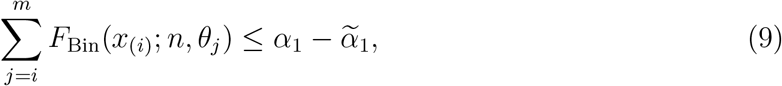

we reject 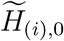 and move to the next joint null that excludes 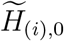; otherwise, we stop and accept the remaining hypotheses. This step-wise procedure asymptotically controls the FWER in testing (3) at 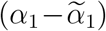 (Hansen, 2005; Romano and Wolf, 2018). Whenever 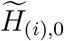 is rejected, the corresponding original hypothesis *H*_(*i*), 0_ is rejected. This procedure is summarized in Step 2 of Algorithm 1.

The step-wise procedure here can be understood as a combination of Holm’s test and Hansen’s adjustment. We can see that, without Hansen’s adjustment, this procedure reduces to Holm’s test. Specifically, the left hand side of (9), with *θ_j_* replaced by *τ_ℓ_*, becomes (*m* – *i* + 1)*p*_(*i*)_, where *p*_(*i*)_ = *F*_Bin_(*x*_(*i*)_; *n*, *τ_ℓ_*) is the *i*th smallest *p*-value among all *p*-values for testing 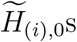. Thus, (9) becomes 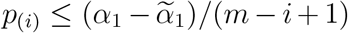, which is Holm’s test. By adjusting for null hypotheses that are “deep in the null”, we have *F*_Bin_(*x*_(*i*)_; *n*, *θ_j_*) ≈ 0 for very large *θ_j_*s and (9) becomes approximately 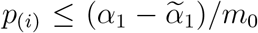, where *m*_0_ is a number that is smaller than *m* – *i* + 1. Therefore, our procedure gains more power than Holm’s test.

### 2.4 A procedure that accepts original hypotheses while controlling the type-II MCER

We can readily modify the procedure in Sections 2.1–2.3 for testing a set of hypotheses that flip the null and alternative hypotheses 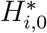 and 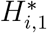 in (2). Thus, a rejection of the new null hypothesis, which is now 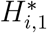, leads to an acceptance of the original hypothesis *H*_*i*,0_. As with the previous procedure, we divide the problem into two sub-problems, the first of which is to construct an at-least-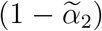-level one-sided CI [0, *τ_u_*] for *τ*\*, where 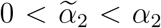, and the second of which is to test the hypotheses that flip the null and alternative hypotheses 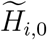 and 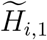 in (3) as well as replace *τ_ℓ_* by *τ_u_*. The details of this procedure are deferred to Appendix and are summarized in Algorithm 2, which can also be found in that section. One important feature of this algorithm is that *τ_u_* is found by using a negative value for *c* in (4), corresponding to the curve *F*_−*c*_(*t*) in Figure 1, guaranteeing that *τ_ℓ_* < *τ_u_*, a result we require in the next subsection.

### 2.5 Combining results of the two procedures

If *H*_*i*,0_ is rejected by the first procedure and non-accepted by the second procedure, it is determined as “rejected”; if *H*_*i*,0_ is accepted by the second procedure and non-rejected by the first procedure, it is determined as “accepted”; if *H*_*i*,0_ is non-rejected and non-accepted, it is determined as “undecided”. Note that our method precludes the possibility that *H*_*i*,0_ is both rejected and accepted, as this would correspond to simultaneously concluding that both 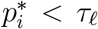 and 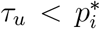 hold simultaneously, which is not possible since *τ_ℓ_* < *τ_u_*. Finally, the overall MCER among the decisions that are either “rejected” or “accepted” (excluding those “undecided”) is controlled at the level that is the sum of the type-I and type-II MCER. The whole workflow of MERIT is depicted in Figure 2.

**Figure 2:**
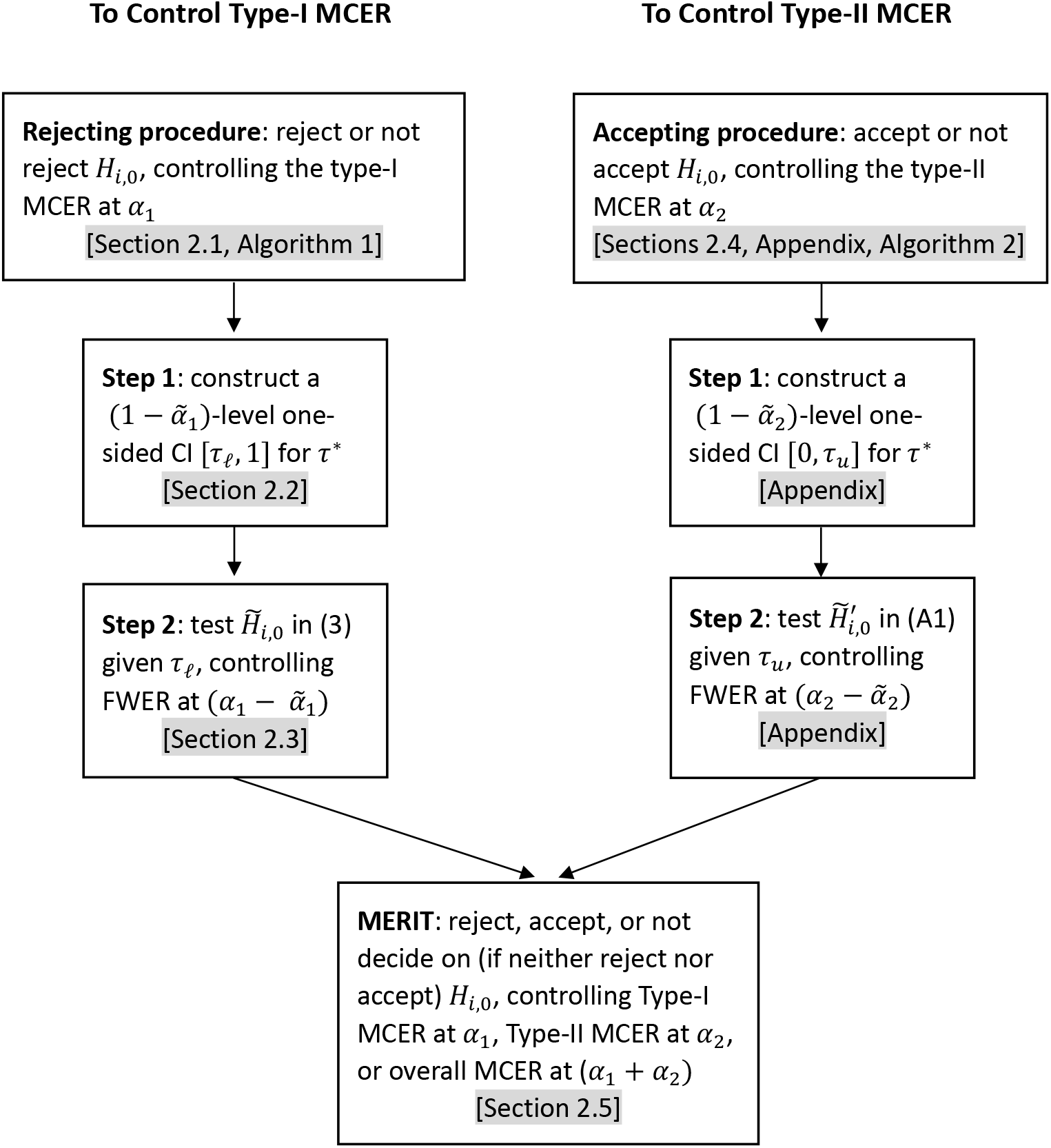
MERIT workflow.

### 2.6 Modifying the GH method for a fixed MC sample size

Because the GH method is fully sequential while MERIT uses a pre-determined MC sample size, it is difficult to compare them directly. For this reason, we created a modified GH method with a fixed stopping rule. We also replaced each two-sided Robbins-Lai interval in GH by two one-sided Wilson intervals (Wilson, 1927) because the Robbins-Lai interval cannot be onesided and because the Wilson interval is more efficient (Brown et al., 2001). In Supplemental Materials S2, we illustrate, using simulated data, that the two-sided Wilson interval (which is the intersection of the two one-sided Wilson intervals) is always narrower than the two-sided Robbins-Lai interval for 0 ≤ *p* ≤ 0.2, the interesting range of *p*-values. To complete our modification of GH, we applied the BH procedure to the upper limits of the Wilson intervals to obtain rejected hypotheses and to the lower limits to obtain accepted hypotheses as in the original GH procedure. Similar arguments to those found in Gandy and Hahn (2014, 2016) can be used to prove our modified GH procedure controls the type-I and type-II MCER.

## 3. Simulation studies

### 3.1 Setup

We conducted extensive simulation studies to evaluate the performance of our method. We evaluated the two procedures of our method (one rejecting hypotheses and one accepting hypotheses). We also compared MERIT to our modified GH method. In addition, we evaluated the naive approach that applies the BH procedure to the MC *p*-values, and we evaluated the type-I and type-II MCER separately. We refer to our method, the modified GH method, and the naive method as MERIT, GH, and NAIVE, respectively.

We considered *m* = 100, 1000, and 5000 tests. We assumed that 80% of *m* tests are under the null and the remaining tests are under the alternative; we sampled the *p*-values for tests under the null independently from the uniform distribution U[0, 1], and sampled the *p*-values for tests under the alternative independently from the Gaussian right-tailed probability model: 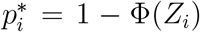, where *Z_i_* ~ N(*β*, 1) and Φ is the standard normal cumulative distribution function. We set *β* to 1.5, 2 and 2.5; a larger value of *β* implies higher sensitivity to reject the alternative hypotheses. For each pair of values of *m* and *β*, we sampled 100 different sets of ideal *p*-values 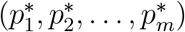. Instead of generating actual data and calculating test statistics, we directly simulated the matrix *I* by independently drawing Bernoulli samples with the “success” rate 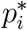. The MC sample size *n* took a wide range of values, i.e., between 5,000 and 1,000,000. The nominal type-I and type-II MCER *α*_1_ and *α*_2_ were both set to 10%; the nominal FDR *ϕ* used to define the BH threshold was set to 10%.

For evaluating the rejecting procedure in each method, we use two metrics, the empirical type-I MCER and sensitivity, the latter of which is

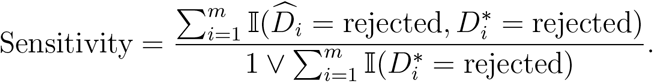

For evaluating the accepting procedure in each method, we also use two metrics, the empirical type-II MCER and specificity, the latter of which is

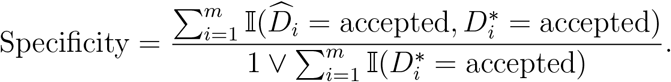

Each result of these metrics was based on 1,000 replicates, which correspond to repeatedly sampled matrix *I* given a set of ideal *p*-values and the MC sample size *n*.

### 3.2 Simulation results

Figure 3 shows the type-I MCER and sensitivity for the three methods, MERIT, GH, and NAIVE, for testing *m* = 1000 hypotheses. Each box displays the variation of results of a method over the 100 sets of ideal *p*-values: the top and bottom lines represent the 75% and 25% quantiles and the middle bar indicates the median. Each point in each box represents results from 1,000 replicates. We can see that, for any combination of *n* and *β*, both MERIT and GH controlled the type-I MCER below the nominal level, while NAIVE did not. MERIT always had higher sensitivity than GH, which is more pronounced when *n* is not very large.

**Figure 3:**
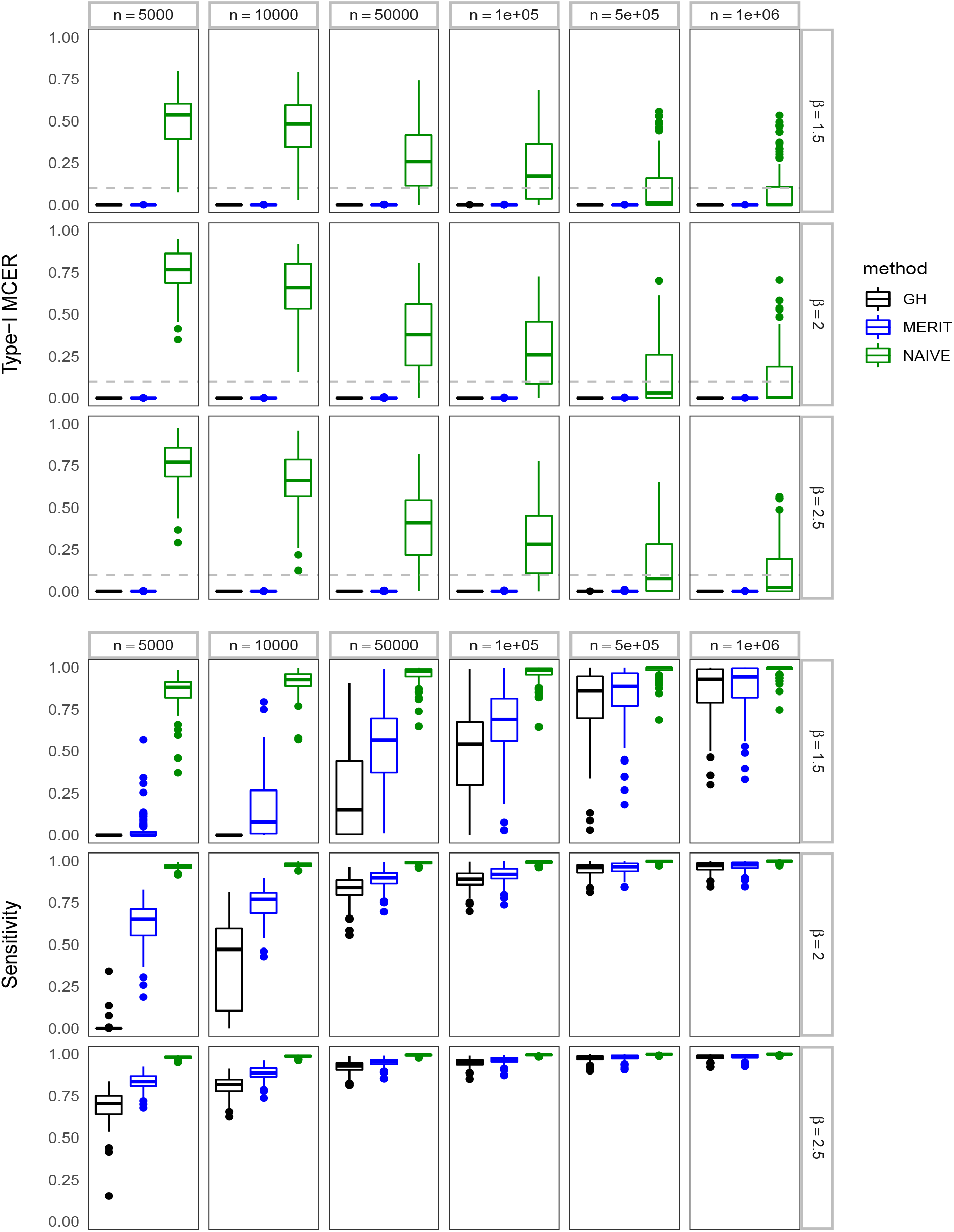
The empirical type-I MCER (upper panel) and sensitivity (lower panel) for *m* = 1000 tests. The gray dashed lines represent the nominal level 10%. Each point represents a different set of ideal *p*-values; results for each set of *p*-values are based on 1,000 MERIT runs.

In fact, MERIT rejected more hypotheses than GH for every set of ideal *p*-values (results not shown). As expected, the type-I MCER of NAIVE was highly inflated when *n* was low; although the error rate decreased as *n* was increased, it was still above the nominal level for many sets of ideal *p*-values even when *n* was as large as one million. One reason that the sensitivity of NAIVE was so high is because it did not allow “undecided” tests, so that every test was categorized either as “rejected” or “accepted”. The results of type-II MCER and specificity are shown in Figure 4, which displays similar patterns as Figure 3. Note that the specificity remained relatively unchanged for different values of *β* because specificity pertains to null hypotheses only.

**Figure 4:**
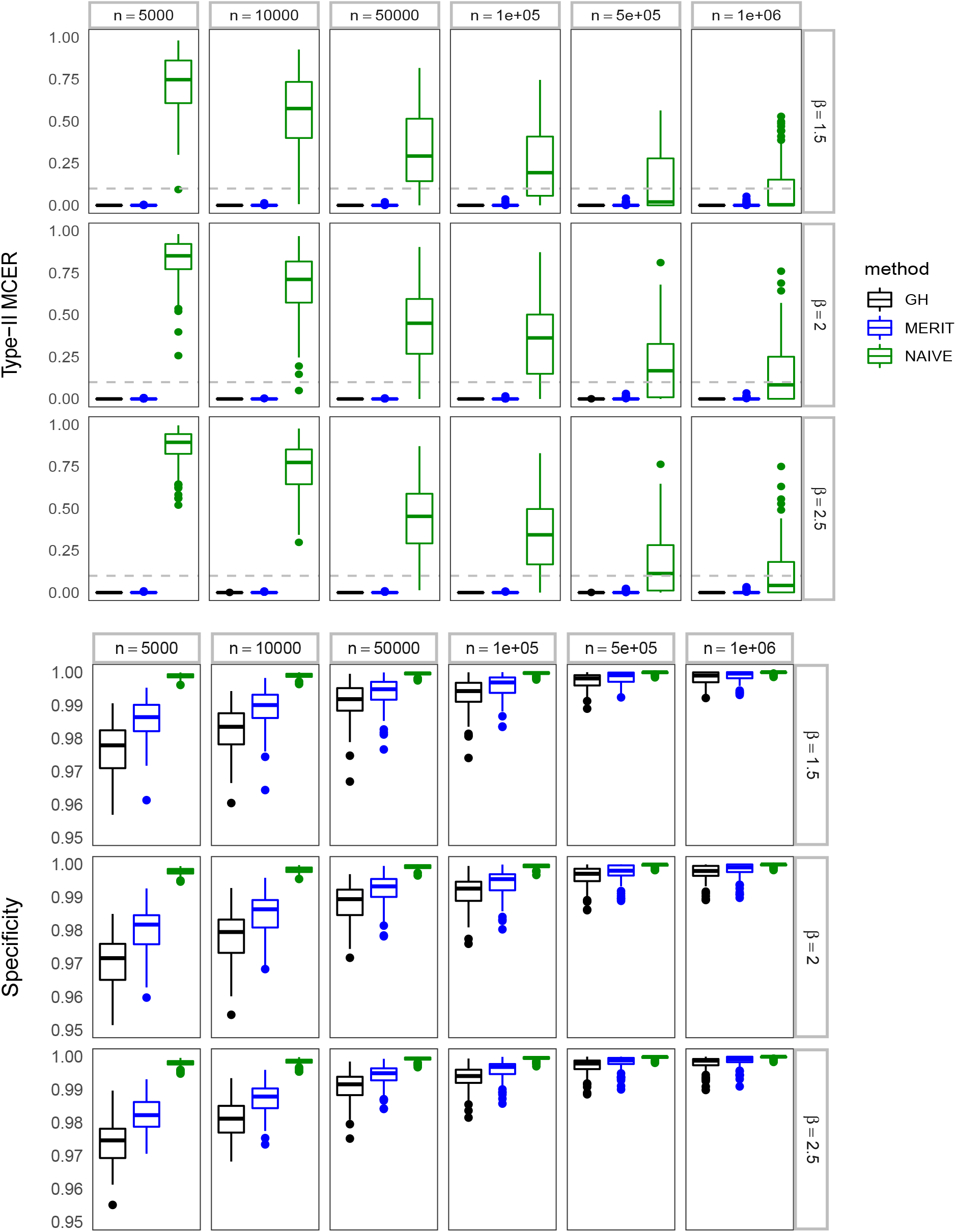
The empirical type-II MCER (upper panel) and specificity (lower panel) for *m* = 1000 tests.

The results for *m* = 100 are shown in Figures S3–S4. With this small number of tests, NAIVE was more likely to control the type-I and type-II MCER because it was less likely to have ideal *p*-values near the threshold of significance. For the same reason, MERIT had higher sensitivity and specificity than in the *m* = 1000 case (for the same values of *n* and *β*). Again, MERIT and GH both controlled the type-I and type-II MCER and MERIT always had higher sensitivity and sensitivity than GH. Note that this behavior with changing *m* is because our ideal *p*-values were assumed to be fairly uniformly spread. If we had assumed a worst-case scenario in which most, or all, ideal *p*-values were at or near the threshold, we would not expect to see this improved behavior when *m* is decreased.

The results for *m* = 5000 are shown in Figures S5–S6. This scale of testing is more commonly seen in modern biological and biomedical studies of omics. Indeed, the prostate cancer data that we will apply our method to has 6033 tests corresponding to 6033 genes. In this case, NAIVE had highly inflated type-I and type-II MCER even with a million MC replicates, implying that the naive results had very poor reproducibility. Again, MERIT and GH both controlled the type-I and type-II MCER and MERIT always had higher sensitivity and sensitivity than GH. Interestingly, while the sensitivity of GH reduced from *m* = 1000 to *m* = 5000, the sensitivity of MERIT stayed quite robust, which suggests an increasing advantage of MERIT over GH for very large numbers of tests.

## 4. Application to the prostate cancer study

We applied MERIT, GH, and NAIVE to detect differentially expressed genes in a prostate cancer study (Singh et al., 2002). The data contains microarray gene expressions for 6,033 genes and 102 subjects comprised of 52 prostate cancer patients and 50 healthy controls. We calculated the *t*-statistic (assuming equal variance) for each gene based on the observed data and *n* MC replicates (by permuting the case-control labels), from which we obtained the matrix *I*. As in the simulation studies, we considered a wide range of values for *n* and we set the nominal type-I and type-II MCER to 10% and the nominal FDR to 10%. For evaluation of reproducibility, we obtained the results for 10 different runs with different seeds for generating MC replicates.

The results are shown in Figure 5. In all cases, MERIT yielded more rejected tests (when there are non-zero rejections) and more accepted tests than GH; the advantage of MERIT is more clear when *n* is between 50,000 and 500,000. Interestingly, the relative numbers of rejections and acceptances of the three methods seem to match well with the sensitivity and specificity in the simulation study with *m* = 5000 and *β* = 1.5. To interpret the results in more detail, we focus on the first run of *n* = 100, 000 MC replicates. MERIT detected 51 genes as differentially expressed and 5961 genes as non-differentially expressed, and left 21 gene “undecided”; in the same run, GH detected 0 gene as differentially expressed and 5954 as non-differentially expressed, and left 79 genes “undecided”. Among the genes detected as differentially expressed by MERIT (or GH), the chance is less than 10% that there exists a gene that would be determined as non-differentially expressed if we obtained an infinite number of MC replicates; we have a similar guarantee for the genes determined as non-differentially expressed.

**Figure 5:**
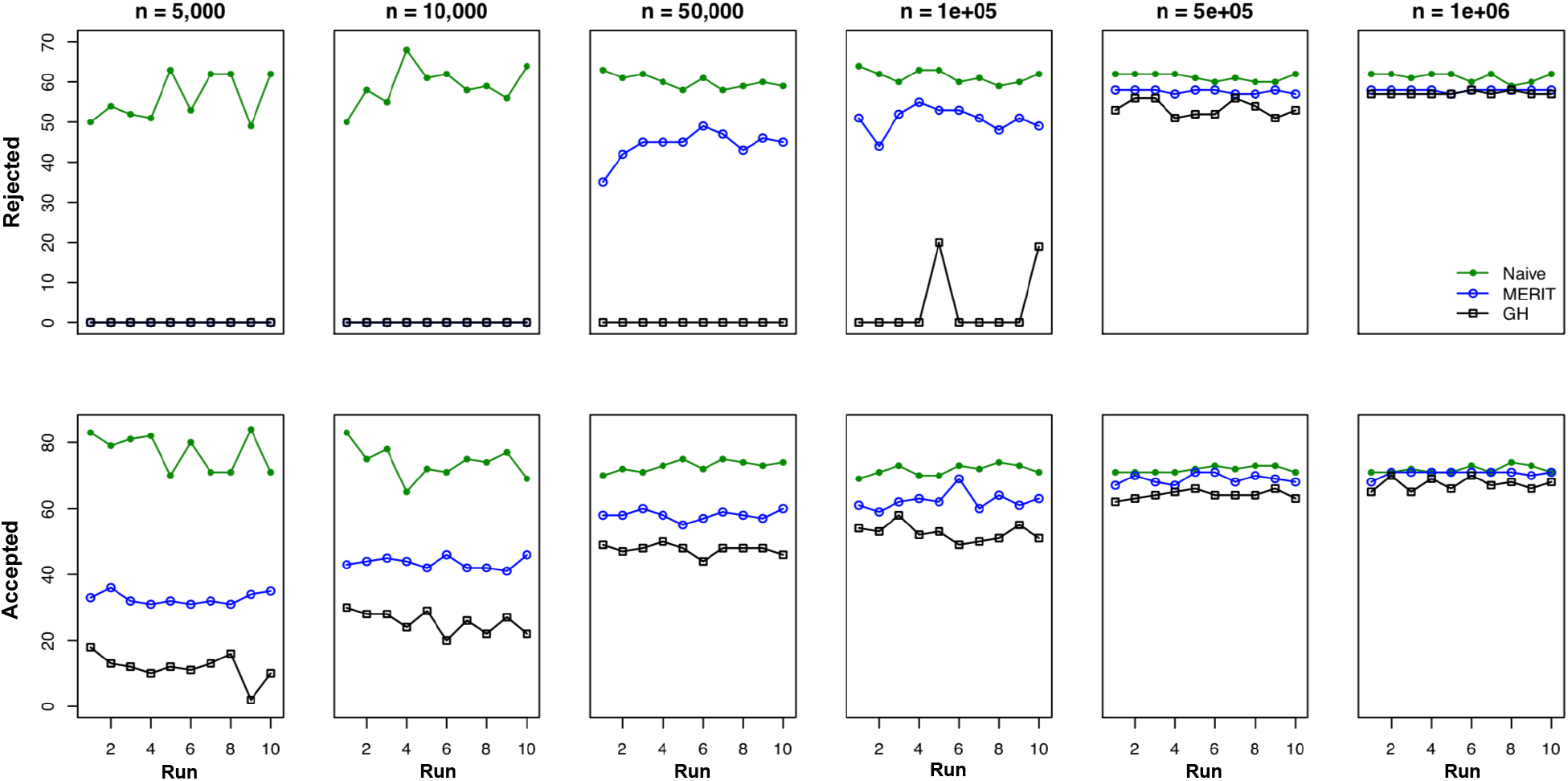
Results from analysis of the prostate cancer data. The upper panel shows the number of rejected hypotheses and the lower panel shows the last two digits (the first two digits are 59; for example, the last two digits 35 correspond to the complete number 5935) of the number of accepted hypotheses. For *n* = 5, 000 and 10,000, both MERIT and GH rejected no hypotheses.

Because MERIT includes a bootstrap step, MERIT will require more run time than NAIVE. For the prostate cancer data, MERIT required ~20 minutes (not including the generation of MC replicates and the calculation of *t* statistic) on a single core for one run of *n* = 1, 000, 000. Because the bootstrap procedure in Step 1 is the most computationally intensive part in our algorithm, the run time can be significantly reduced by paralleling the computation of bootstrap replicates to multiple cores. For example, one run of MERIT for the prostate cancer data with *n* = 1, 000, 000 required only ~2 minutes on 10 cores.

## 5. Discussion

Monte-Carlo error is a threat to reproducibility in two related ways, when testing hypotheses using resampling techniques such as permutation or bootstrap. First, MC error may lead to false rejection of a hypothesis that we would not reject given the ideal *p*-value. Second, if we re-run an MC hypothesis testing procedure, we may obtain a different list of rejected or accepted hypotheses. Note that when the outcome “undecided” is available, these two types of MC error are distinct. The control of MCER guaranteed by MERIT means that, even if two runs give different lists of rejected or accepted hypotheses, we are assured that the chance of reaching the wrong conclusion about any hypothesis is bounded by level *α*. Thus, while successive runs of MERIT may have different lists of rejected or accepted hypotheses, these lists will only differ by switches between “rejected” and “undecided”, or between “accepted” and “undecided”, with probability at least (1 — *α*). The situation is very different for the NAIVE procedure where the BH threshold is applied directly to the MC *p*-values. Here the absence of an “undecided” category means that *all* errors are switches between “rejected” and “accepted”, even though the overall (experiment-level) false discovery rate may be controlled. Many of these false discoveries will be due to MC error alone.

MERIT achieves control of type-I MCER at level *α*_1_ by lowering the BH threshold for rejecting hypotheses. We note that we could address the chance of switches between “rejected” and “undecided” hypotheses by lowering this threshold even further. If we lowered the threshold such that the probability of observing any switches between “rejected” and “undecided” was bounded by *α*_3_, then we could control the probability of observing *any* irreproducible rejections by *α*_1_ + *α*_3_. This control would, however, come at the cost that some hypotheses that can be rejected by controlling type-I MCER at level *α*_1_ will no longer be rejected by the new criterion. Still, it may be worth considering. Finally, the number of ideal *p*-values near the threshold of significance controls the number of “undecided” hypotheses in MERIT, and the distance between the MC *p*-value and this threshold presumably controls the likelihood of a switch between “rejected” and “undecided”. In our experience, when there is a large number of “undecided” hypotheses, changing the nominal FDR level *ϕ* to a value where there are relatively few empirical *p*-values can sometimes reduce the number of “undecided” hypotheses.

When developing MERIT, we chose to control the family-wise MCER, rather than a percentage or rate of MC errors among all decided tests, since this MC “false discovery rate” may be misleading. The chance of an MC error for hypotheses with ideal *p*-values well below the BH threshold is typically fairly small; the presence of these “easily rejected” hypotheses in a false-discovery-like list of MC errors can therefore lead to fairly loose control of MC error for hypotheses that are closer to the BH threshold, even though the overall error rate is controlled. In contrast, controlling the family-wise MCER bounds the probability of *any* MC error, which seems to be more appropriate in the current context.

MERIT (as well as our modification of GH) differ from the original GH proposal in that type-I and type-II MCER can be separately controlled. This may be useful for some users who care more about discoveries (i.e., rejections) than they care about lists of hypotheses for which the null hypothesis is accepted. For these users, type-I MCER may be the only error worth controlling. In this situation, we may set the type-II MCER to *α*_2_ = 0 so that we only obtain two possible outcomes: “rejected” or “undecided”. Alternatively, a small type-II MCER *α*_2_ can be selected so that hypotheses that are very far from the threshold of significance can still be labeled as “accepted”.

Here we have considered only MC schemes that have a fixed stopping criterion. We feel this is reasonable, as our impression is that most large-scale MC hypothesis testing reported in the biological sciences used a fixed MC sample size. This also enables us to achieve higher efficiency, as it allows for a fairly sophisticated correction for multiple testing to be applied only once, after all tests have been completed. For fairness, we have also modified the original, sequential GH approach to allow for a more efficient approach when a fixed stopping criterion is used. Even with this boost, we showed that MERIT outperformed GH unless the MC sample size was 1,000,000 or greater, after which performance of the two methods was similar. In this sense, our method is the best available approach to MC hypothesis testing that features control of MCER with a fixed stopping criterion. We have implemented our method in the R package MERIT available on GitHub at XXX.

## Appendix A procedure that accepts original hypotheses while controlling the type-II MCER

We consider the following hypotheses that “flip” the hypotheses in (2):

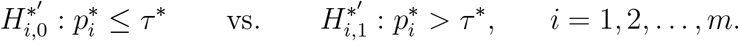

Note that a rejection of 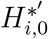 corresponds to an acceptance of the original hypothesis *H*_*i*,0_. Recall that a type-II MC error occurs when 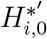 is true (hence 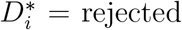) but rejected using the MC *p*-value (hence 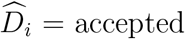). Then, a test procedure based on MC *p*-values that controls the FWER for testing 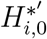 at *α*_2_ would control the type-II MCER at *α*_2_. Here, we develop a two-step procedure following the same idea as in Sections 2.1–2.3. In the first step, we construct an at least 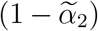-level one-sided CI (0, *τ_u_*] for *τ*\*, where 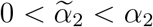. In the second step, we consider the revised hypotheses, treating *τ_u_* as fixed:

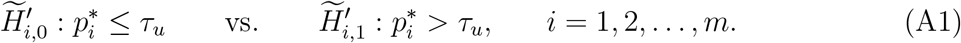

We develop a test procedure that tests (A1) while controlling the FWER at level 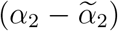.

Then, by the relationship that

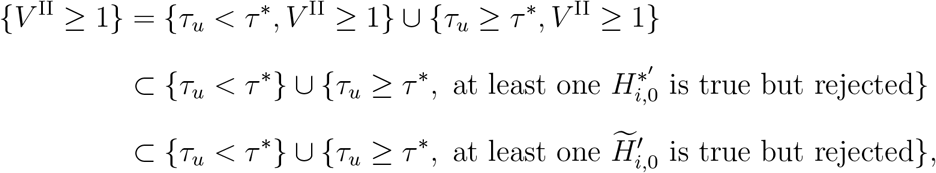

the type-II MCER Pr(*V*^II^ ≥ 1) is always bounded by the sum of Pr(*τ_u_* < *τ*\*) and the FWER in testing (A1), which is *α*_2_. As the default choice, we choose 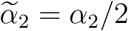.

### Step 1: construct a 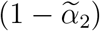-level one-sided CI [0, *τ_u_*] for *τ*\*

We find *τ_u_* from a set of *p*-values that are “upper limits” for the ideal *p*-values. To this aim, we consider the following sets of *p*-values, with each set indexed by a positive continuous value *c*:

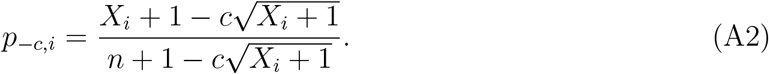

Note that we impose the restriction that 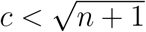; otherwise 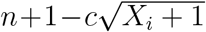 may become 0 or even negative and then *p*_−*c,i*_ is not well-defined. However, negative values of the numerator *are* allowed, as illustrated in Figure 1 where the curve *F*_−*c*_(*t*) has non-zero value at *t* = 0. As *c* goes to 0, *p*_−*c,i*_ converges to 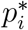. When *c* is sufficiently large, *p*_−*c,i*_ is asymptotically smaller than 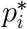, as least for small 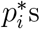.

Let *F*_−*c*_(*t*) be the EDF of (*p*_−*c*,1_,…, *p*_−*c,m*_) and *τ*_−*c*_ be the BH cutoff. If *F*_−*c*_(*t*) is always no less than *F*\*(*t*) over (0, *ϕ*], as shown in Figure 1 (left), we must have *τ*_−*c*_ ≥ *τ*\*. It follows that

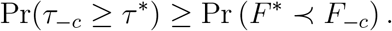

To construct a 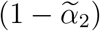-level one-sided CI (0, *τ_u_*] for *τ*\*, we are interested in *c* that satisfies

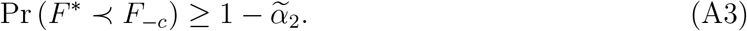

The probability on the left of (A3) increases as *c* increases (because the decrease of *p*_−*c,i*_ leads to the increase of *F*_−*c*_(*t*) at a given *t*). Thus we wish to find the smallest *c* that satisfies (A3), denoted by *c_u_*. Finally, we set the upper limit *τ_u_* to *τ_c_u__*.

We obtain *c_u_* using the same bootstrap replicates as in Section 2.2. Define the bootstrap *p*-values indexed by *c* as

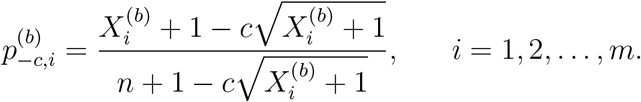

Let 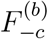 be the EDF for 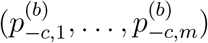. We wish to find *c_u_* to be the smallest *c* that satisfies

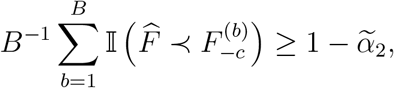

which is the empirical version of (A3). To this aim, we find, for every *b* = 1,…, *B*, the smallest *c* that guarantees 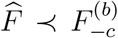, denoted by 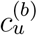, and then set *c_u_* to be the 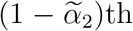 quantile of 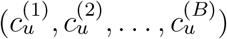. The procedure here is summarized in Step 1 of Algorithm 2.

### Step 2: test hypotheses in (A1) given *τ_u_* while controlling the FWER at 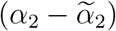

Let 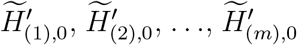 be the ordered hypotheses (A1) that correspond to the ordered observed test statistics *x*_(1)_, *x*_(2)_,…, *x*_(*m)*_. We start by testing the global null 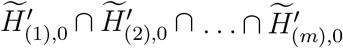, which means that all 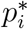 are less than *τ_u_*. We use *x*_(*m*)_ as the test statistic for this test. Following Romano and Wolf, we propose to calculate the *p*-value under the global null:

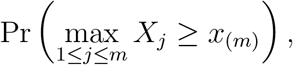

where each *X_j_* is a binomial random variable with *n* trials and “success” rate 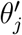 that is under the null. Using the Bonferroni inequality, we obtain an upper bound of the *p*-value to be

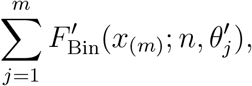

where 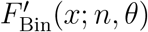 denotes Pr(*X* ≥ *x*) for *X* ~ Bin(n, *θ*). Using Hansen’s adjustments for null hypotheses that are “deep in the null”, we choose 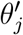 to be

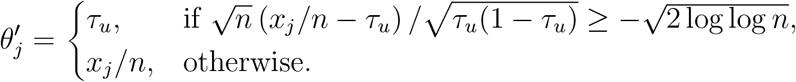

Using this 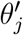, if

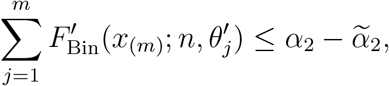

we reject 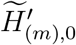 and move to the next joint null 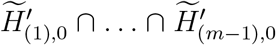 that excludes 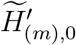; otherwise, we stop and declare that none of the hypotheses should be rejected. In general, for testing the joint null 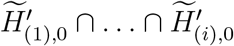 for any *i*, we use *x*_(*i*)_ as the test statistic. If

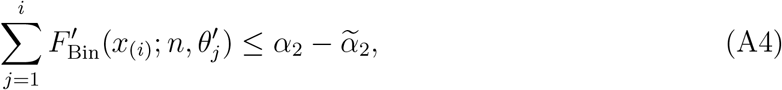

we reject 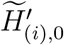 and move to the next joint null that excludes 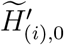; otherwise, we stop and accept the remaining hypotheses. This step-wise procedure asymptotically controls the FWER in testing (A1) at 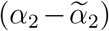. Whenever 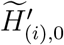 is rejected, the corresponding original hypothesis *H*_(*i*) 0_ is accepted. This procedure is summarized in Step 2 of Algorithm 2.

#### Algorithm 2 The procedure that accepts original hypotheses while controlling type-II MCER

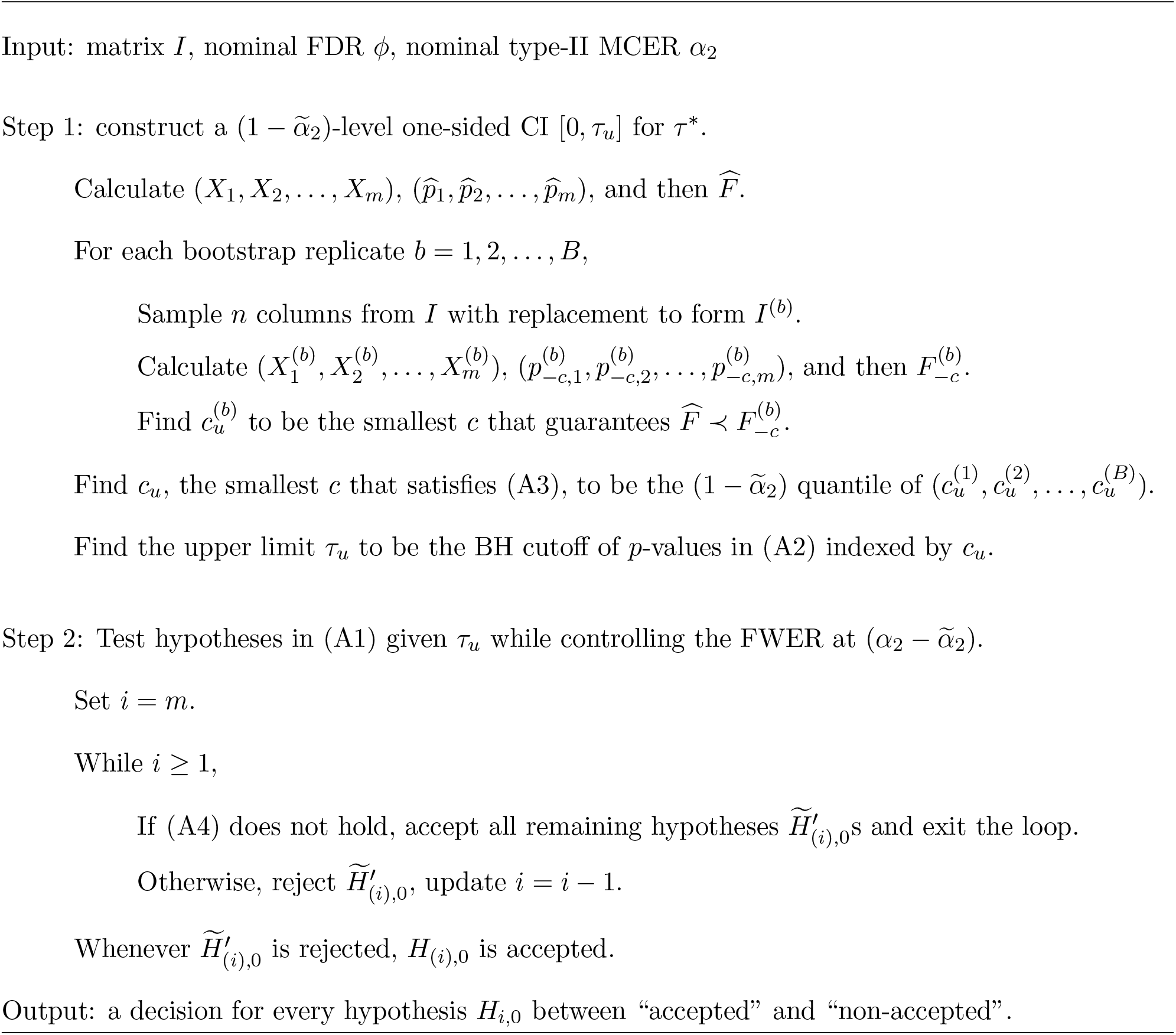

## Supplementary Materials

### S1. Comparing different schemes of splitting the nominal type-I MCER between the two sub-problems

We used simulations to compare different schemes for splitting the nominal type-I MCER in terms of the sensitivity of rejecting alternative hypotheses. Here, the nominal type-I MCER (*α*_1_) was set to 10%. We considered 5 different schemes, 1:9, 3:7, 5:5, 7:3, and 9:1, where, for example, 1:9 means that the error rate 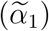 assigned to the first sub-problem was 1% and the error rate 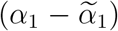 assigned to the second sub-problem was 9%. We considered *m* = 1000 tests and drew 100 sets of ideal *p*-values. All results in Figure S1 suggest that the 5:5 scheme is a robust choice.

**Figure S1:**
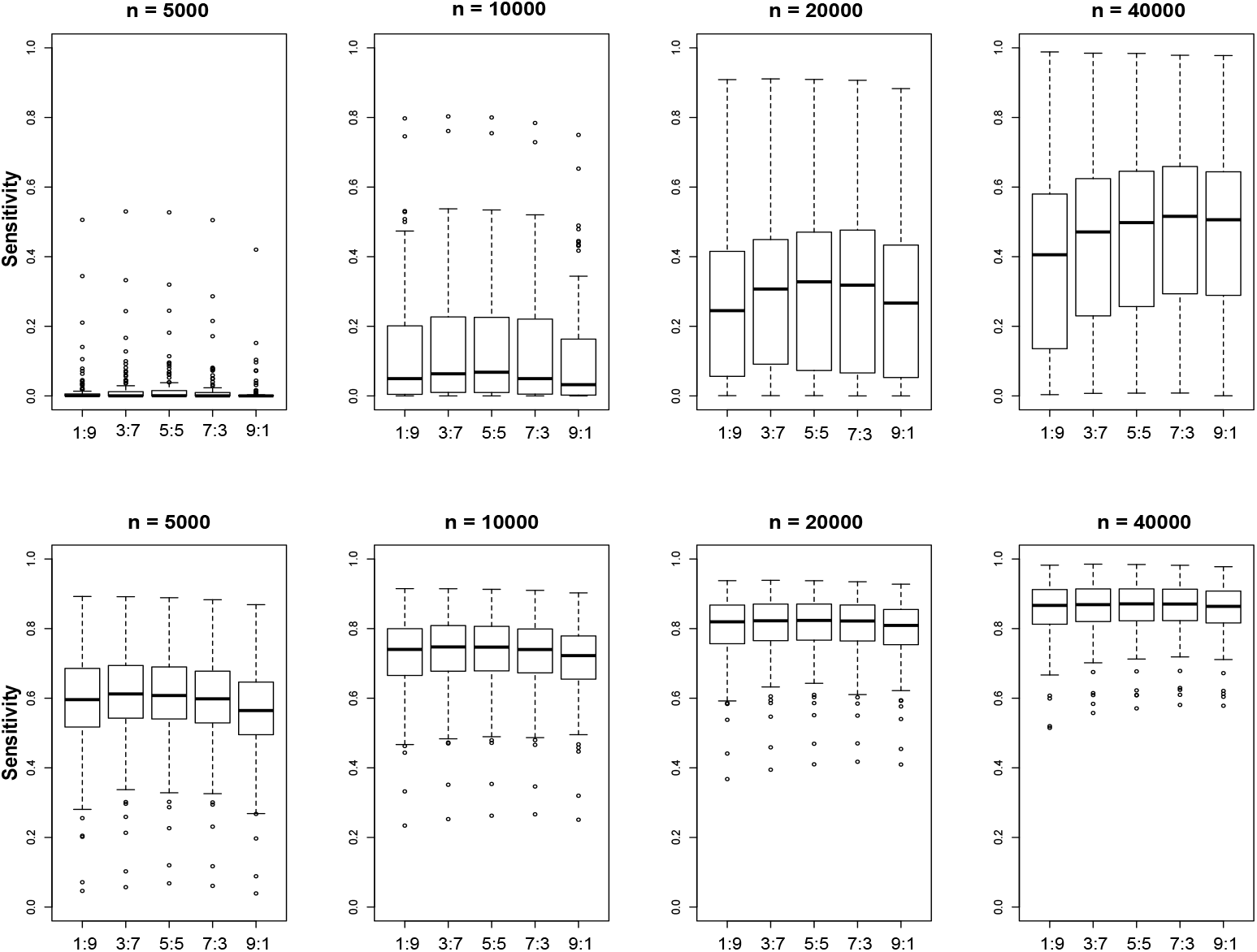
The empirical sensitivity of MERIT using different schemes for splitting the nominal type-I MCER. The upper and lower panels are based on *β* = 1.5 and 2.0, respectively.

### S2. Comparing the Robbins-Lai and Wilson intervals

We used simulations to compare the width of the two-sided Robbins-Lai interval and the two-sided Wilson interval. For each *p* ~ [0, 0.2] (as large values of *p* are irrelevant because they will not lead to rejection of hypotheses for any reasonable error rate), we simulated the total number of exceedances *X* ~ Bin(*n*, *p*), where *n* = 100, 000. Then we calculated the width of the two intervals and obtained the average width over 1000 replicates. Finally, we formed the ratio of the average widths (Robbins-Lai:Wilson) and displayed it in Figure S2. We can see that the ratio is always above 1 (between 1.6 and 2).

**Figure S2:**
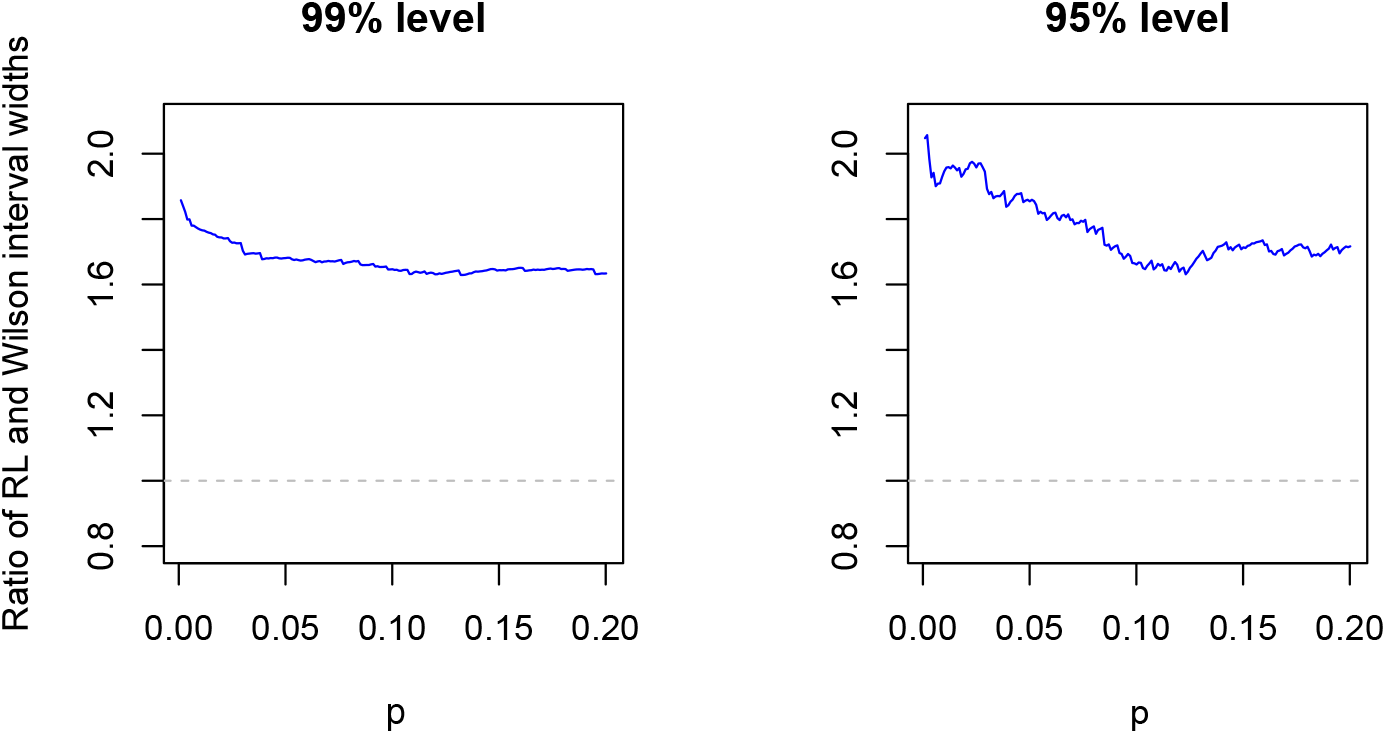
The ratio of the widths of the Robbins-Lai and Wilson intervals. The left and right panels pertain to 99%- and 95%-level intervals, respectively.

**Figure S3:**
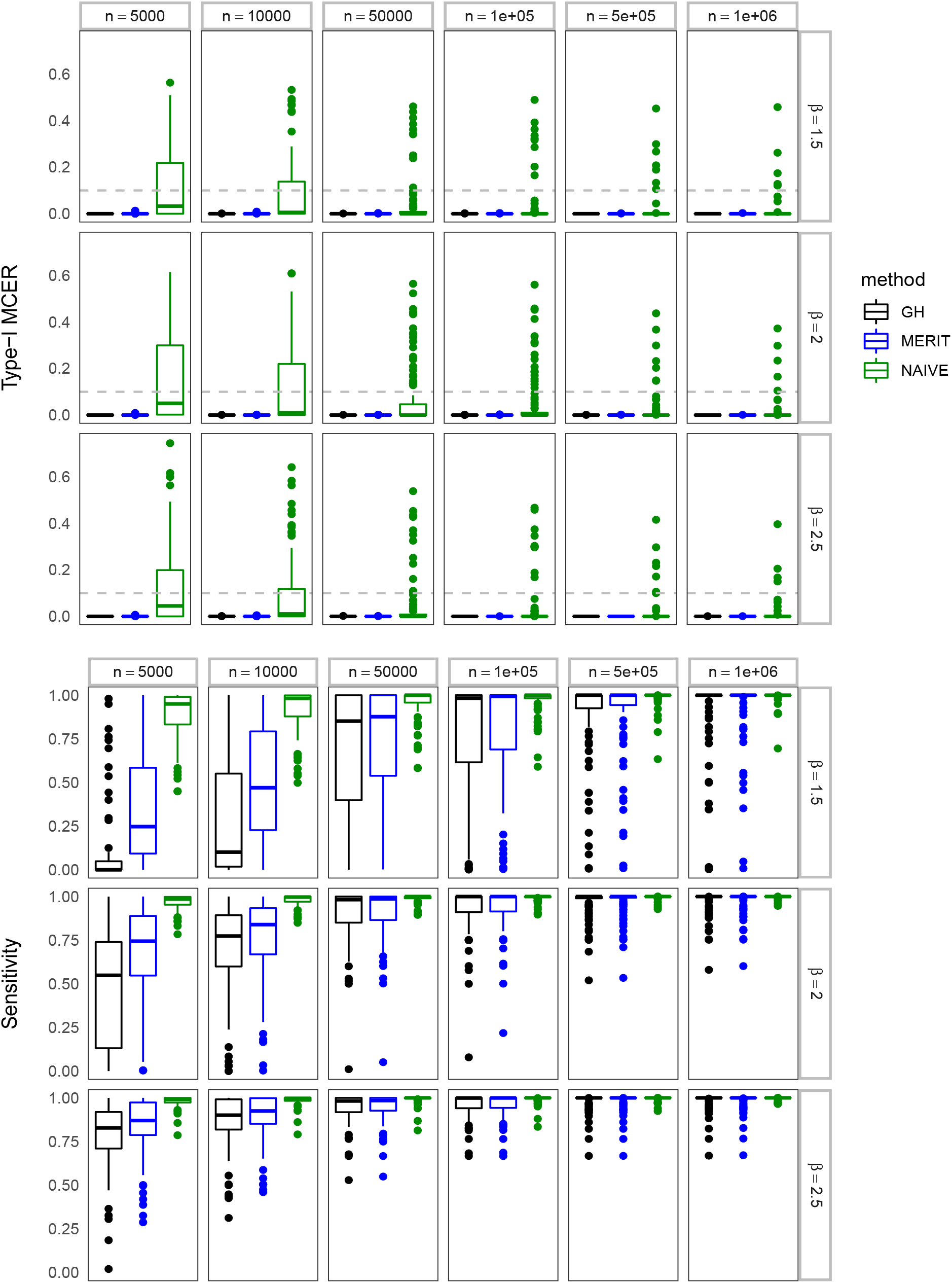
The empirical type-I MCER (upper panel) and sensitivity (lower panel) for *m* = 100 tests.

**Figure S4:**
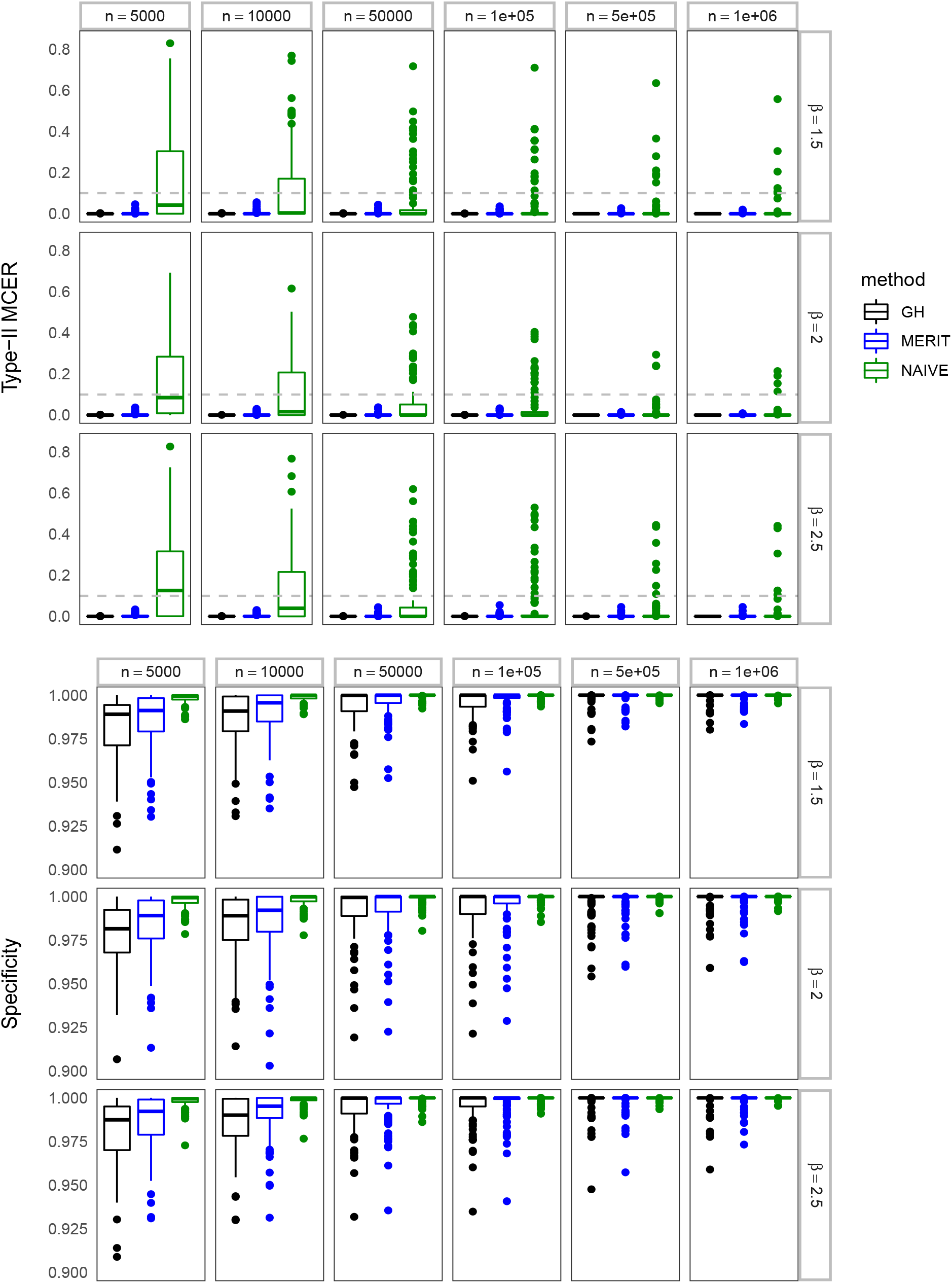
The empirical type-II MCER (upper panel) and specificity (lower panel) for *m* = 100 tests.

**Figure S5:**
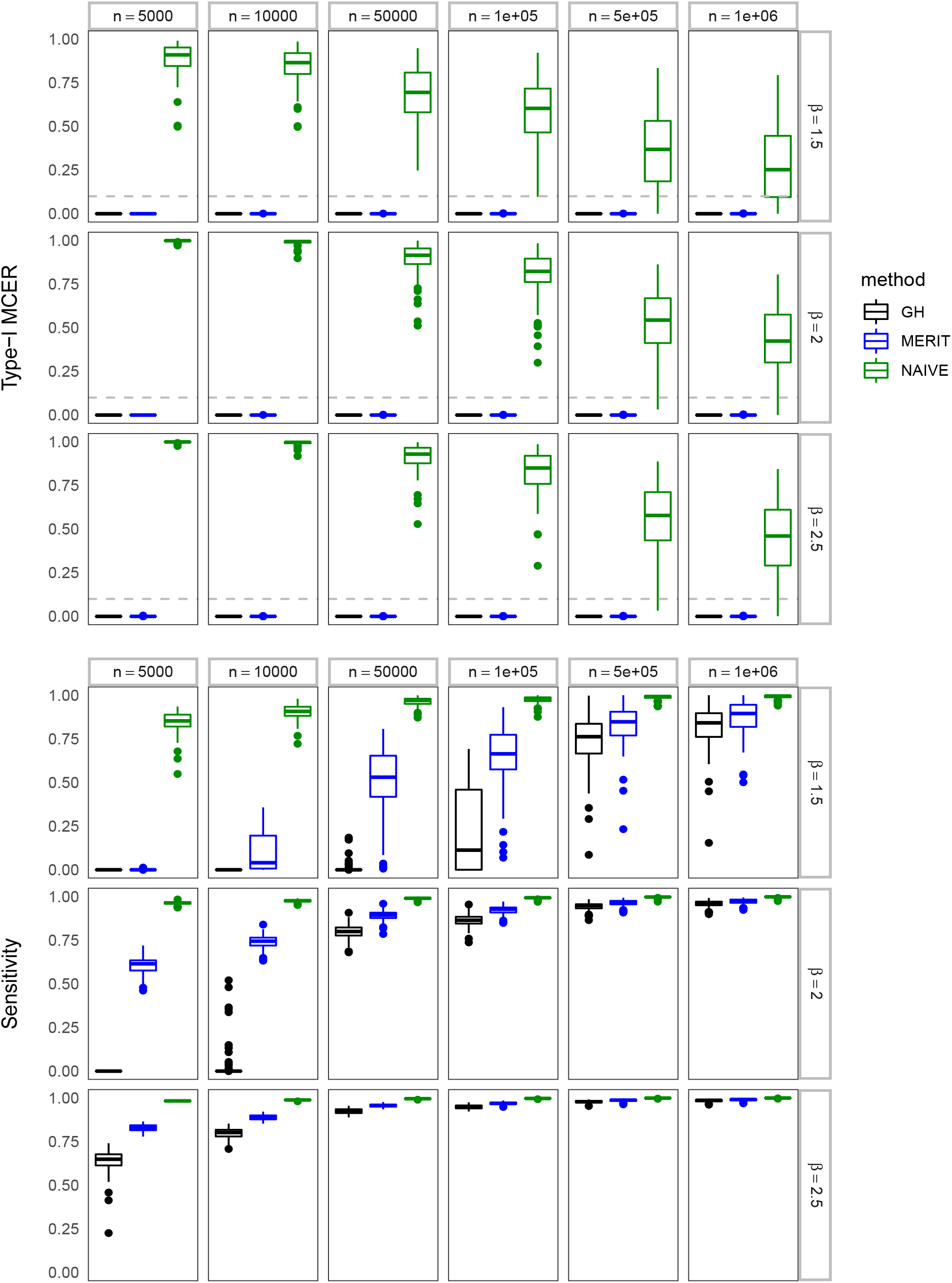
The empirical type-I MCER (upper panel) and sensitivity (lower panel) for *m* = 5000 tests.

**Figure S6:**
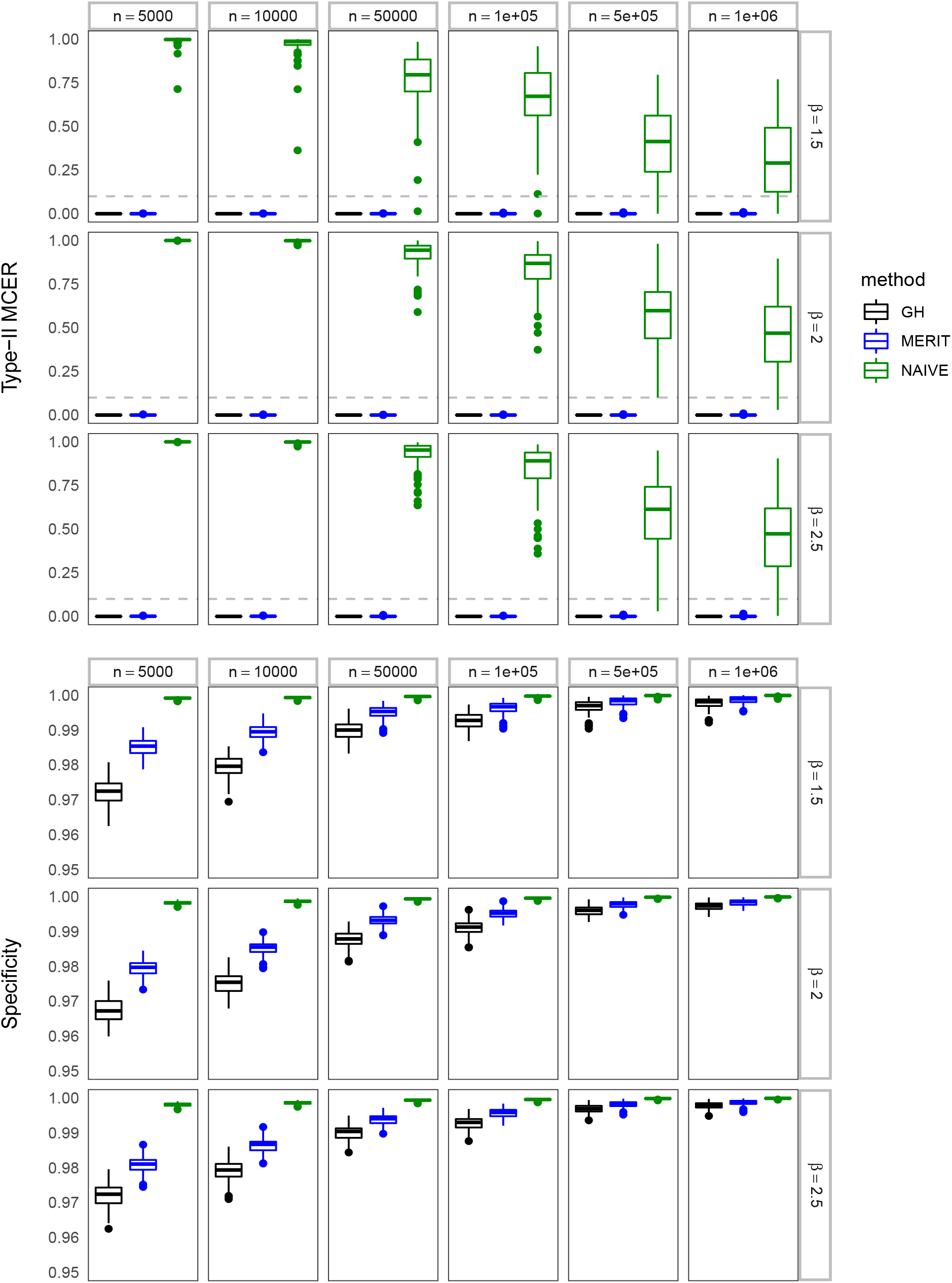
The empirical type-II MCER (upper panel) and specificity (lower panel) for *m* = 5000 tests.

